# Injury stimulates stem cells to resist radiation-induced apoptosis

**DOI:** 10.1101/688168

**Authors:** Divya A Shiroor, Tisha E Bohr, Carolyn E Adler

## Abstract

Stem cells are continuously exposed to multiple stresses including radiation and tissue injury. Because stem cells are central drivers of tissue repair and regeneration, it is essential to understand how their behavior is influenced by these stressors. Planarians have an abundant population of stem cells that are rapidly eliminated after radiation exposure via apoptosis. Low doses of radiation eliminate the majority of these stem cells, allowing a few to remain [1]. Here, we combine radiation with injury to define how surviving stem cells respond to tissue damage. We find that injuries induce stem cells to persist, but only if injured within a defined window of time surrounding radiation, and only immediately adjacent to the wound. Stem cells persist for several days without any proliferation. Instead, they are retained near the wound due to suppression of apoptosis, which we quantify in stem cells by combining FACS with Annexin V staining. Tissue injury is known to induce apoptosis in differentiated cells [2], and we hypothesize that these dying cells confer apoptosis resistance onto nearby stem cells. Indeed, pharmacological induction of cell death with cycloheximide is sufficient to prolong survival of radiated stem cells even in the absence of injury. Together, our results suggest a model in which dying cells provide a transient protective signal to nearby stem cells, altering their susceptibility to radiation-induced apoptosis.

## Results and discussion

Planarians are a highly tractable model system to study regeneration, due to a large heterogeneous stem cell population consisting of pluripotent cells and organ-specific progenitors [3–5]. Upon injury, stem cells proliferate to initiate regeneration [6,7]. Injuries trigger a vast wound-induced transcriptional program [6,8,9], and rampant apoptosis occurs at the wound site, but how these changes impact stem cell behaviors remains unclear. However, the sheer number of stem cells obscures visualization of cell behavior. Here, we overcome this problem by exposing animals to radiation, which causes loss of most or all of the stem cells, in a dose-dependent manner. For example, a lethal dose of 6000 rads or higher eliminates all stem cells via apoptosis [2], and consequently inhibits regeneration [10,11]. However, at sublethal doses, some cells survive, eventually repopulating the animal [1,12], providing an opportunity to track stem cell behavior after injury and radiation.

### Stem cells persist adjacent to wounds

To establish a context to study stem cell behavior, we exposed animals to sublethal doses of radiation, which eliminates all but a few stem cells [1]. We used a dose of 2000 rads because it preserved a quantifiable population of stem cells after exposure. To observe the kinetics of stem cell loss after radiation, we first assessed the rate of stem cell depletion after exposure to 2000 rads. We measured expression of *smedwi-1*, an Argonaute family protein whose transcript is exclusively expressed in stem cells [11,13]. In accordance with prior studies [12,14], we observed a pronounced decrease in *smedwi-1* transcript levels 1 day after exposure (Figure 1A). Previous results have shown that decapitation 5 days after radiation failed to increase stem cell numbers [1]. Because so many stem cells are lost by this late timepoint, we investigated whether injuries inflicted while stem cells are still present would impact their behavior. Therefore, we decapitated animals soon after radiation and monitored stem cell numbers with *in situ* hybridization for *smedwi-1* two days later (Figure 1B). If decapitation occurred within 24 hours of radiation, or immediately after exposure, stem cells were retained adjacent to the wound. This result indicates that stem cells can respond to injury after radiation, but only if the injury occurs within a critical window.

**Figure 1:**
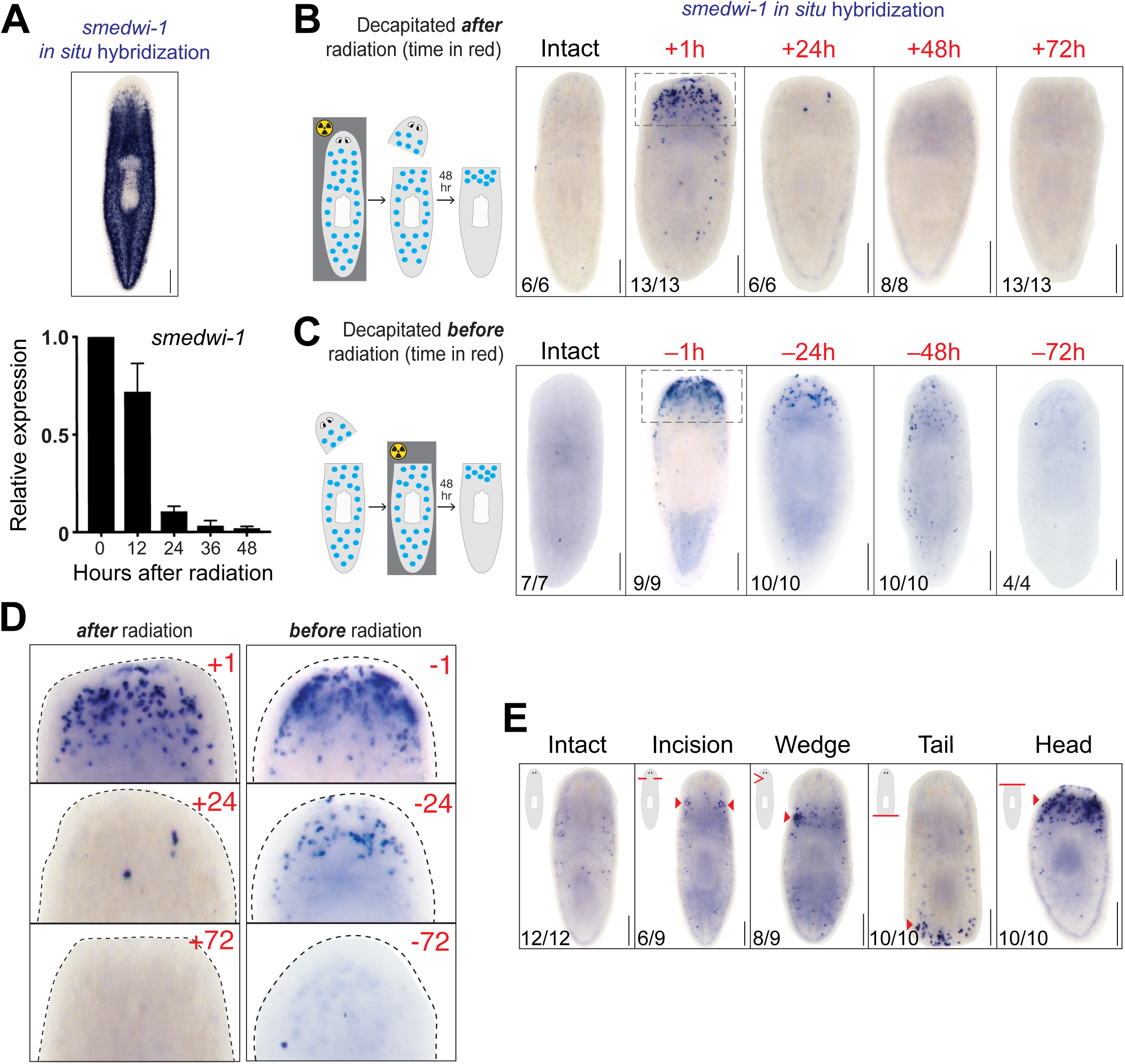
Injury within a ‘critical window’ of radiation induces local stem cell persistence. A) Top, *smedwi-1 in situ* hybridization in an unirradiated animal. Bottom, quantitative RT-PCR of *smedwi-1* relative to GAPDH at specified times after radiation exposure. Error bars=SD. (B-E) All panels show *smedwi-1 in situ* hybridization in animals fixed 48 hours after injury. Animals were exposed to 2000 rads. B) Animals were exposed to radiation, then decapitated afterwards at indicated times (red). C) Animals were decapitated at indicated times prior to radiation (red). D) Magnified images of blastemas from boxed regions in B and C, decapitated at indicated times before or after radiation. E) Injuries inflicted as indicated in top left corner by red lines. Animals were injured within one hour of radiation exposure. Arrowheads indicate stem cells. Scale bars = 250µm.

To define the full duration of this response, we reversed the order of radiation and decapitation, and performed amputations prior to radiation exposure. In a mirror image of the sensitivity after radiation, we find that animals amputated 24 hours prior to radiation also retain stem cells near the wound (Figure 1C). If decapitation occurs 3 days before or after radiation, stem cells fail to respond (Figure 1D). Therefore, timing of decapitation relative to radiation is crucial. Decapitation inflicted prior to radiation generates a transient signal that lasts for 24 hours, while injury after radiation indicates that stem cells are only receptive to this signal for up to 24 hours. Given the robust and consistent presence of stem cells when we decapitated animals within an hour of radiation exposure (n>1000 animals over multiple experiments), we used this paradigm for all further experiments unless otherwise specified.

Our results suggest that upon injury, stem cells are maintained adjacent to the wound site. To examine the dependence of this response on either the position or size of the injury, we varied the location and severity of the wound. Regardless of the type or position of the injury (tail removal or wedges removed pre-pharyngeally), stem cells were retained locally, adjacent to the wound (Figure 1E). In addition, incisions on either side of the head that did not remove any tissue also resulted in stem cell persistence. Together, these data demonstrate that stem cells perdure locally, near any injury site.

To verify that the cells present around the wound were bonafide stem cells, we evaluated other known canonical, stem-cell-specific transcripts [14–16]. We examined the co-expression of *smedwi-2* together with *smedwi-1* using double fluorescent *in situ* hybridization (FISH) [11,17]. All *smedwi-1* cells also expressed *smedwi-2*, confirming that *smedwi-1* identifies stem cells (Supplementary Figure 1A). Other markers of the stem cell population include Histone *H2b* [16], the RNA-binding protein *bruli* [18], and *soxp-1* [17,19,20]. Decapitation induced maintenance of all of these markers adjacent to the wound site (Supplementary Figure 1B), indicating that the phenotypes we observe are not unique to *smedwi-1*.

In addition, planarian stem cells are a heterogeneous population, consisting of both pluripotent stem cells capable of restoring all of the animal’s organs [21], and lineage-restricted progenitors. To determine if injury causes persistence of these distinct populations in radiated animals, we examined the expression of *tgs-1* and *zfp-1*, a marker for epidermally-fated stem cells [17,22]. Both of these stem cell markers were present at the wound site (Supplementary Figure 1B), demonstrating that pluripotent and lineage-restricted progenitor stem cells respond to injury after radiation.

### Stem cell persistence is not due to proliferation

Injury in planarians causes stem cells to enter the cell cycle in two waves. A body-wide increase in proliferation occurs 6 hours after any kind of wound, and a second proliferative burst occurs locally 48 hours after tissue removal [6,23]. Because we observed such a sharp increase in stem cell number after radiation and amputation compared to uninjured controls, we reasoned that proliferation might contribute to this increase. To test this, we stained animals with an antibody recognizing phosphorylated histone H3 at serine10 (H3P) 2 and 7 days after radiation and decapitation. Unirradiated control animals had expected amounts of proliferation at both timepoints. However, we did not detect any proliferating cells in radiated decapitated animals, despite the pronounced presence of *smedwi-1*^*+*^ stem cells adjacent to the wound (Figure 2A). Therefore, proliferation does not contribute to the stem cell persistence we observe after radiation and decapitation.

**Figure 2:**
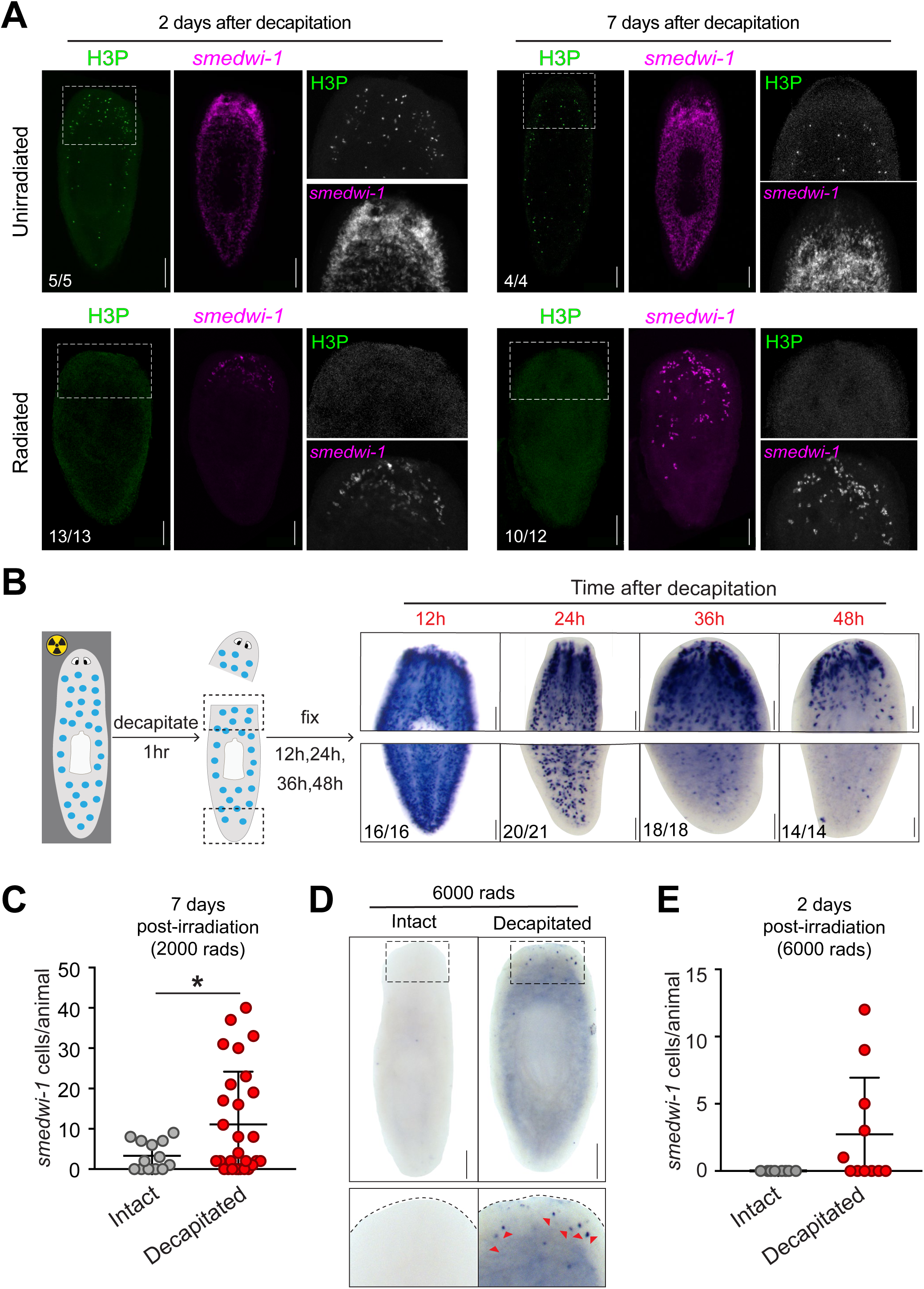
Stem cell persistence is not due to proliferation. A) Phosphohistone H3 (H3P) antibody and *smedwi-1* expression 2 and 7 days after decapitation in unirradiated (top) or radiated (bottom) animals. Dashed boxes highlight zoomed in regions on the right. B) Decapitated animals fixed at indicated timepoints. Boxed regions of *smedwi-1* expression are shown on the right. C) Quantification of *smedwi-1*^*+*^ stem cells in animals 7 days after radiation. Asterisk = p<0.003, student’s t-test. Circles represent individual animals. D) Animals exposed to lethal radiation (6000 rads) and decapitated within an hour. Red arrows indicate *smedwi-1*^*+*^ cells. Second row of images are zoomed regions indicated by box. E) Quantification of *smedwi-1*^*+*^ stem cells in D. Circles represent individual animals. For all experiments (except D and E), animals were exposed to 2000 rads. Scale bars: A, B=100µm, D=250µm.

If proliferation does not generate new stem cells after injury, an alternative possibility is that stem cells in injured animals survive radiation. To address this possibility, we visualized early events after radiation by fixing animals at 12 hour intervals after decapitation. At 12 and 24 hours, *smedwi-1* cells were still present throughout the body of decapitated animals. However, at later timepoints, we observed a strong depletion in the tail, distal to the injury (Figure 2B), while being maintained near the wound and persisting for up to 7 days after decapitation (Figure 2C). These data indicate that instead of stimulating proliferation, injury retains radiated stem cells around the wound site. To further test this, we increased the radiation dose from sublethal (2000 rads) to lethal (6000 rads), known to eliminate 100% of stem cells [11,14,19]. We reasoned that if injury induces stem cells to persist, we would detect them after lethal radiation. As expected, 2 days after lethal radiation, intact animals had no stem cells present. By contrast, decapitated animals retained some stem cells around the wound site (Figure 2D and 2E). Injury, therefore, facilitates stem cell persistence after radiation, regardless of the dose.

The prolonged survival of stem cells induced by decapitation suggested that these cells may retain the capacity to differentiate. We therefore examined differentiation with neuronal markers restricted to the head. *Cintillo* and *ovo* are exclusively expressed in small populations of sensory and photoreceptor neurons, respectively [24,25]. Two weeks after radiation exposure and decapitation, we observed partial restoration of both *cintillo* and *ovo* (Supplemental Figure 2A), demonstrating that radiated stem cells are capable of normal differentiation.

We next examined whether persisting radiated stem cells could contribute to long-term survival of the animal. Because intact animals died within 18-28 days after exposure to higher doses (>2000 rads), we also included lower doses where animals survived indefinitely (Supplementary Figure 2B). All animals decapitated after exposure to 500 rads survived. However, doses of 1000 rads or higher caused decapitated animals to die earlier than their intact counterparts (Supplementary Figure 2B). This indicates that beyond a certain threshold, radiation impairs the ability of stem cells to facilitate regeneration, likely because of sustained DNA damage.

### Stem cells resist radiation-induced apoptosis

Radiation exposure causes loss of stem cells via apoptosis [2]. Our results therefore suggest that injury may alter the propensity of radiated stem cells to undergo apoptosis. To test this possibility, we used fluorescence-activated cell sorting (FACS), a well-established strategy for isolating a highly enriched population of stem cells (X1) based on DNA content [11,26]. We combined FACS with Annexin V staining, which is routinely used to label apoptosis in intact cells [27,28], to directly measure cell death occurring exclusively in stem cells.

To validate this Annexin V-FACS method in planarians, we first tested whether radiation exposure would increase rates of apoptosis in stem cells. As expected, we observed a precipitous decline in the X1 population from 12.7% in unirradiated animals to 1.6% and 0.66% 1 and 2 days after radiation exposure, respectively (Figure 3A). This decline in the X1 population was accompanied by an increase in the percentage of Annexin V-positive stem cells. Within the stem cell population, we found that in unirradiated controls, only 6% of stem cells were Annexin V-positive, but stem cells from radiated animals displayed significantly higher proportions of X1 cells with increased staining intensity (33.1% and 69.9% 1 and 2 days after radiation) (Figure 3B and 3C). This expected decline in stem cells following radiation, accompanied by an increase in Annexin V staining, confirms that this technique can be used to identify stem cells undergoing apoptosis.

**Figure 3:**
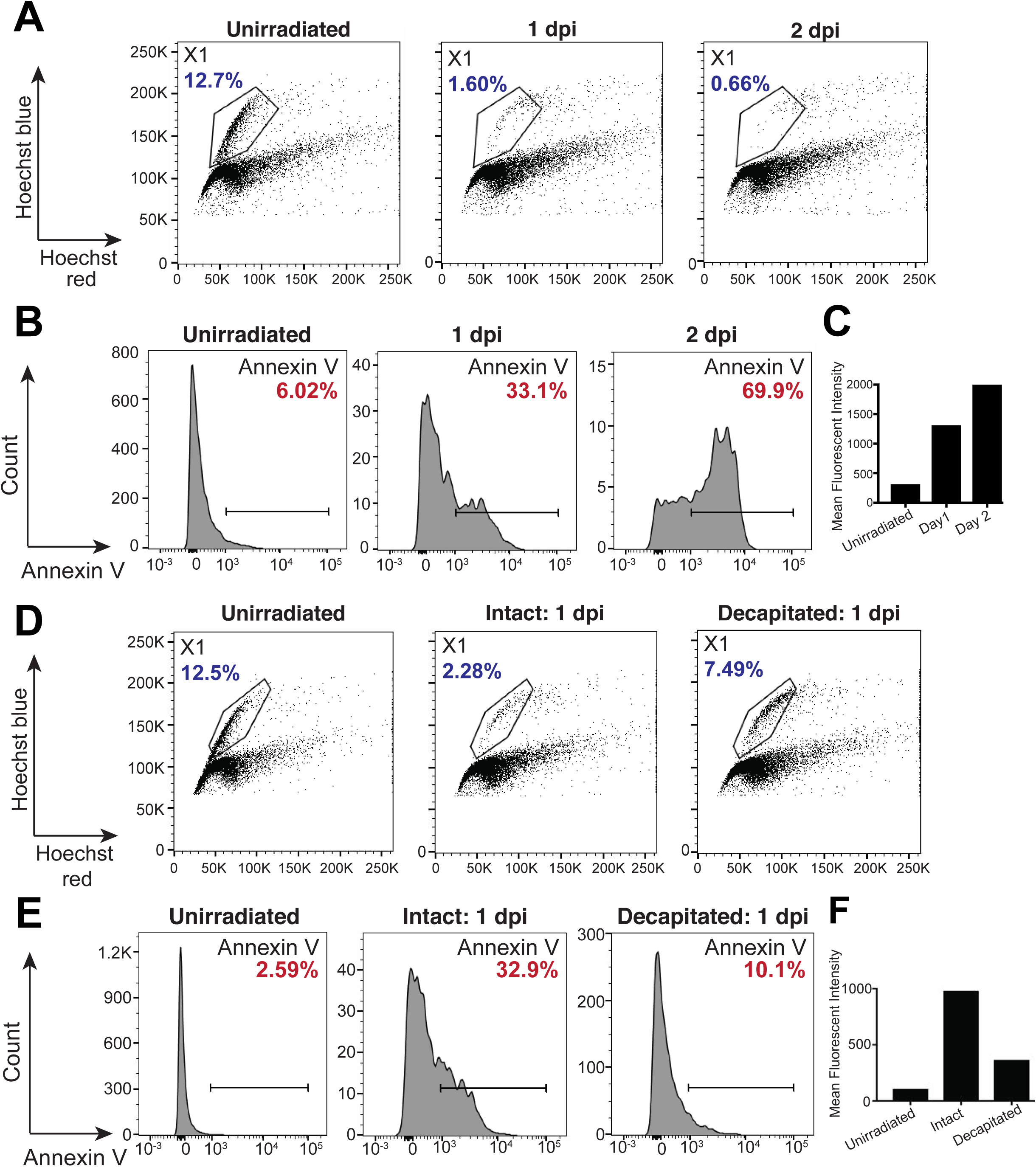
Stem cells resist radiation-induced apoptosis after injury. A) FACS plots of Hoechst-stained X1 cells, 1 or 2 days after radiation (dpi). X1 gate is encircled with black lines. Percentage of stem cells within the total population is represented in blue. B) Histograms of X1 cells from A stained with Annexin V. The percentage of stem cells with Annexin V staining above 10^3^ is shown in red, and highlighted by brackets. C) Graph of mean fluorescent intensity of Annexin V (from one representative experiment) calculated from bracketed region in B. D) FACS plots of Hoechst-stained X1 cells. Animals were decapitated within 1 hour of radiation, and processed 24 hours later. X1 gate is encircled with black lines. Percentage of stem cells within the total population is represented in blue. E) Histograms of X1 cells from D stained with Annexin V. The percentage of stem cells with Annexin V staining above 10^3^ is shown in red, and highlighted by brackets. F) Graph of mean fluorescent intensity of Annexin V (from one representative experiment) calculated from bracketed region in D. Figures are representative of N>2 experiments. All X1 cells were isolated from the pre-pharyngeal region 24 hours after exposure to 2000 rads.

If injury is in fact providing an escape from radiation-induced apoptosis, we predicted that radiated, decapitated animals would have fewer Annexin V-positive stem cells as compared to their intact counterparts. Using Annexin V-FACS, we compared the abundance of apoptotic stem cells in radiated intact and decapitated animals. Consistent with our *in situ* hybridization results, 1 day after radiation, decapitated animals had more X1 cells as compared to intact animals (7.49% decapitated versus 2.28% intact) (Figure 3D). This population of cells also showed less Annexin V staining than intact animals, both in cell numbers and intensity (32.9% in intact animals versus 10.1% in decapitated animals) (Figure 3E and 3F), indicating that injury interrupts the initiation of apoptosis. Given that we find fewer cells initiating apoptosis in radiated, decapitated animals, we suggest that tissue injury confers resistance to radiation-induced apoptosis.

### Death of differentiated cells is sufficient to promote stem cell survival

In planarians, wounding induces immediate transcriptional programs [8,29] and migration of stem cells toward the injury [30]. We hypothesized that these early wound response genes might be involved in local persistence or accumulation of stem cells after decapitation. We knocked down three of these genes (*junl, fos-1*, and *jnk*) prior to radiation and decapitation [29,31,32] and found no difference in stem cell numbers (Supplementary Figure 3). Similarly, knockdown of *snail* [33] did not impact stem cell numbers. Therefore, we concluded that migration and early wound response genes were unlikely to account for the stem cell maintenance we observe.

In planarians, injury induces a high concentration of apoptotic cells locally at the wound site [2]. We combined whole-mount TUNEL (terminal deoxynucleotidyl transferase-mediated dUTP nick end labeling) with *in situ* hybridization for *smedwi-1* to assess the proximity of stem cells to dying cells, and found that they are adjacent, but do not coincide (Figure 4A). Studies in *Drosophila*, Hydra and zebrafish have shown that dying cells influence adjacent cells. This occurs either by inducing proliferation [34–37], or by altering cell survival and fate in a non-autonomous manner [38–40]. Because we confirmed that no proliferation occurs (Figure 2A), we instead considered the possibility that injury-induced apoptosis may also promote stem cell survival.

**Figure 4:**
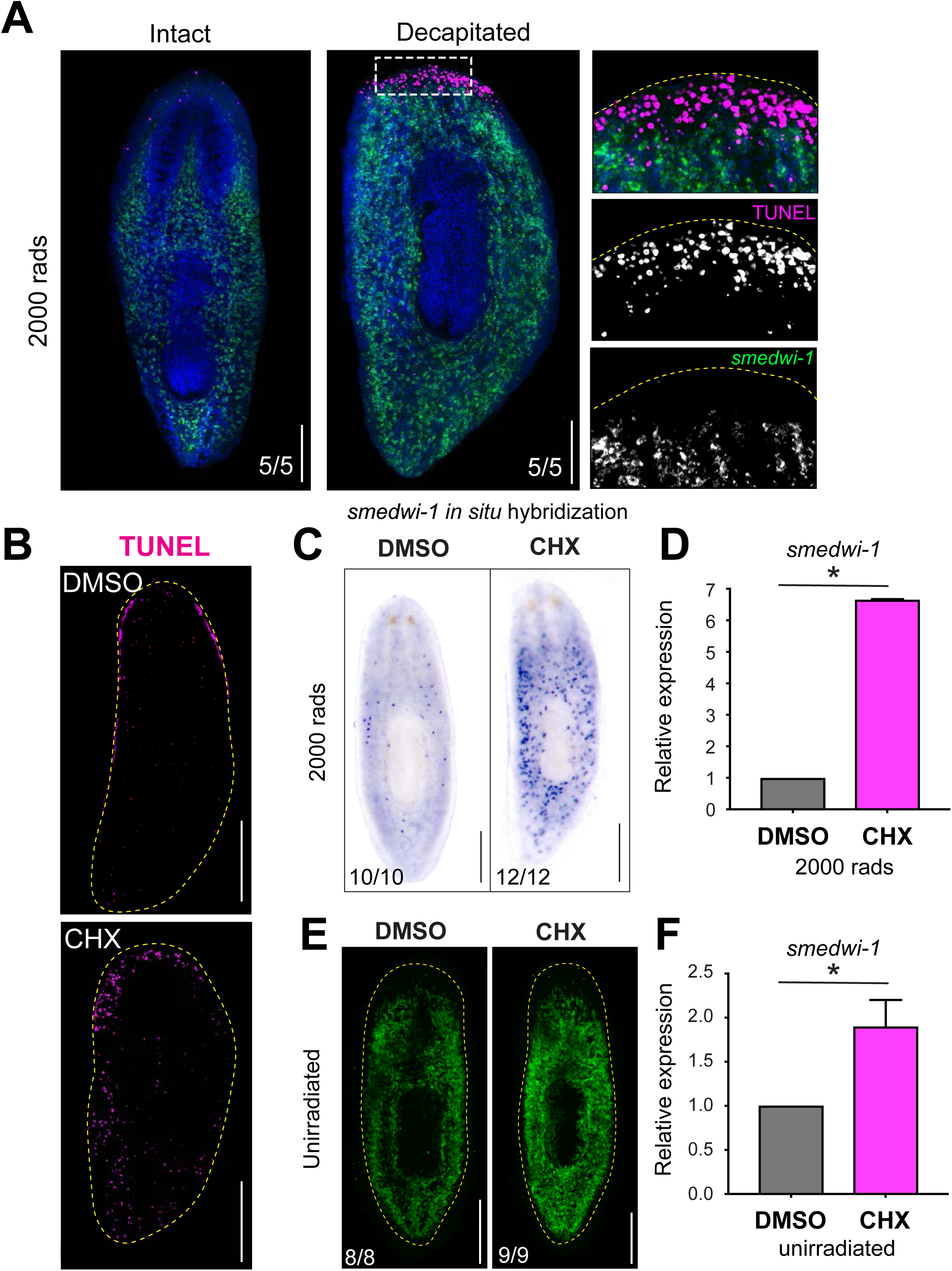
Apoptosis promotes stem cell survival. A) Confocal projections of animals stained for TUNEL (magenta) and *smedwi-1 in situ* hybridization (green), 4 hours after radiation and decapitation. Dashed box highlights zoomed regions on the right. B) Confocal projections of animals stained with TUNEL, after soaking in DMSO (control) or cycloheximide (CHX) for 3 days. C) *smedwi-1 in situ* hybridization of animals soaked in DMSO or CHX for 2 days prior and 1 day after radiation exposure, then fixed 1 day later. D) Quantitative RT-PCR of *smedwi-1* relative to GAPDH in animals soaked and radiated as in C. Asterisk represents p<0.0014, Welch’s t test. Error bars=SD. E) Confocal projections of fluorescent *smedwi-1 in situ* hybridization in animals soaked in DMSO or CHX for 3 days, then fixed 1 day later. F) Quantitative RT-PCR of *smedwi-1* relative to GAPDH in animals soaked as in E. Asterisk represents p<0.003, Welch’s t test. Error bars=SD. In A, C, and D, radiation dose was 2000 rads. Scale bars: A =100µm. B,C,E = 250µm.

To directly test whether dying cells may prolong stem cell survival after radiation, we took advantage of previously published pharmacological agents. The translation inhibitor cycloheximide is known to induce apoptosis due to loss of rapidly turned over anti-apoptotic proteins such as Mcl-1 [41]. Similarly, planarians soaked in cycloheximide exhibit elevated levels of TUNEL staining (Figure 4B) [2]. To test whether increased apoptosis was sufficient to induce stem cell survival, we soaked animals in cycloheximide for 2 days prior to radiation exposure, and one day afterward. Cycloheximide-soaked animals had markedly increased stem cell numbers after radiation as compared to controls (Figure 4C). We confirmed this increase by measuring *smedwi-1* transcript levels with quantitative RT-PCR (Figure 4D). These results suggest that apoptosis alone, even in the absence of any mechanical injury, is sufficient to prolong stem cell survival after radiation.

Because radiation itself induces apoptosis of stem cells, and they do not survive in intact animals, we ruled out the possibility that they could provide a protective effect to neighboring stem cells. Instead, we hypothesize that the protective effect must originate from other cell types that die. Notably, we also observed a slight but clear increase in stem cell abundance following cycloheximide exposure in unirradiated animals (Figure 4E and 4F). These data imply that dying cells can also regulate stem cell behaviors during homeostasis, even without the added stress of radiation. Our data strongly suggest that dying cells influence stem cell survival and prevalence, but do not rule out potential contributions by other mechanisms such as dedifferentiation.

## Conclusion

The role of apoptosis in planarian homeostasis and regeneration has been mysterious. Ours is the first study to provide evidence that dying cells can influence stem cell behavior in these animals. Using radiation, which causes stem cells to rapidly initiate apoptosis, we uncovered a new component of the stem cell response to injury. By pairing injury with radiation, we find that stem cells evade entering apoptosis, but only if the injury occurs within a ‘critical period’ surrounding radiation. Evidence from other model organisms has shown non-autonomous effects linking apoptotic cells to stem cell behaviors [38,42–44]. However, our approach affords a unique perspective because of the acuteness with which the injury is applied, and the direct outcomes on adult stem cells. The pronounced concentration of stem cells around the wound suggests that injury induces local apoptosis, which in turn enables stem cell survival within a defined radius of the wound. Our results indicate that apoptosis may impact stem cells both during homeostasis and regeneration, outside the context of radiation. This work provides a paradigm for further dissection of the mechanisms used by stem cells to interpret and respond to diverse environmental or wound-induced stresses.

## Acknowledgements

We thank Jason Pellettieri and John Dustin for generously providing us with their optimized TUNEL-FISH protocol; Jason Pellettieri, Nicole Lindsay-Mosher and Nicolas Buchon for comments on the manuscript; Chris Donahue at the Cornell University Flow Cytometry Core for assistance in optimizing Annexin V staining protocols; Sabrina Solouki and Avery August for use of their Cell Counter; and Robert Weiss for maintaining the irradiator. D.A.S. was supported by a GRA Fellowship from the Cornell College of Veterinary Medicine. This work was supported by Cornell University startup funds and a Seed Grant from the Cornell University Stem Cell Program to C.E.A.

## Author Contributions

D.A.S. conceived, designed and conducted experiments; analyzed data; wrote the manuscript.

T.E.B. conducted experiments; analyzed data; edited the manuscript.

C.E.A. conceived and designed experiments; wrote the manuscript.

## Figure legends

Supplementary Figure 1: Stem cells localizing to the wound site express stem-cell-specific transcripts.

A) Double fluorescent *in situ* hybridization of *smedwi-1* (green) and *smedwi-2* (red). Imaged region is highlighted by the box in the cartoon. Percentage represents *smedwi-1*^*+*^ cells that co-express *smedwi-2* (white arrowheads). 800 cells were quantified from n=6 animals. Scale bar = 15µm.

B) *In situ* hybridization for stem cell markers indicated. Scale bar = 250µm.

All animals were decapitated within an hour of radiation and fixed 2 days after exposure to 2000 rads. Boxed region represents zoomed areas in the bottom row.

Supplementary Figure 2: Radiated stem cells differentiate but do not enhance animal survival.

A) *In situ* hybridization for markers of photoreceptors (*ovo*) and sensory neurons (*cintillo*) at 0 or 14 days after decapitation, with and without irradiation. Dashed boxes highlight zoomed regions on the bottom. Radiation dose=2000 rads. Scale bar = 250µm.

B) Kaplan-Meier curves of animal survival of intact (top) and decapitated (bottom) animals after radiation. Radiation doses indicated in graph. n=30 animals.

Supplementary Figure 3: Wound response and migration candidates are not required for stem cell persistence after radiation.

*In situ* hybridization of *smedwi-1* in radiated intact (top) and amputated (bottom) animals subjected to RNAi as indicated. Scale bar = 250µm. Boxed region represents zoomed areas in the bottom row.

## STAR Methods

### Planarian care and irradiation

Asexual planarians from the *Schmidtea mediterranea* clonal line CIW4 were kept in Montjuïc salts at 20°C [45]. Animals used for experiments were between 1-5mm in length and starved for 5-7 days. Planarians were irradiated on a J.L. Shepherd & Associates Mark I-68 Irradiator, and dosage was calculated based on exposure time.

### qRT-PCR

A total of 10 animals were processed per biological replicate with 3 biological replicates and 3 technical replicates per condition. Animals were collected in Trizol (Thermo Fisher 15596018) in Lysing Matrix D Tubes (MP Biomedicals 116913100) and homogenized using a Bead Bug microtube homogenizer (Benchmark). RNA was extracted according to the Trizol protocol. cDNA was synthesized using Superscript™ VILO™ (Life Technologies 11754250). PCR mixes were made using TaqMan™ Gene Expression Master Mix (Life Technologies). Custom primers are available at ThermoFisher sequences for *smedwi-1* (AI89MBJ) and GAPDH (AI6RPY3). PCR was run on an Applied Biosystems Viia7 Real Time PCR System and quantified using Ct methods.

### Fixations

Animals were fixed and labeled as previously described [46]. Briefly, animals were killed using 7.5% N-acetyl-cysteine in PBS for 10 minutes and fixed in 4% paraformaldehyde for 30 minutes at room temperature. After fixation, worms were rinsed twice with PBSTx (PBS + 0.3% Triton X-100). PBSTx was replaced with pre-warmed reduction solution (PBS+ 1% NP-40+ 50mM DTT + 0.5% SDS) and animals were incubated at 37°C for 10 minutes. After rinsing twice with PBSTx, animals were dehydrated in a methanol series and stored at −20°C.

### Whole-mount *in situ* hybridizations

Colorimetric *in situ* hybridizations were performed as described in [46] and fluorescent *in situ* hybridizations as in [47] with minor modifications. Briefly, animals fixed as above were rehydrated, bleached (5% Formamide, 1.2% hydrogen peroxide in 0.5x SSC) and treated with proteinase K (4 µg/ml in 1x PBSTx, Thermo Fisher 25530049). Following overnight hybridizations at 56°C, samples were washed 2x each in wash hybe (5 min), 1:1 wash hyb:2X SSC-0.1% Tween 20 (10 min), and 2X SSC (30 min), 0.2X SSC (30 min) at 56°C followed by 3 x 10 minute MABT washes at room temperature. Subsequently, animals were placed in blocking solution (0.5% Roche Western Blocking Reagent and 5% inactivated horse serum diluted in MABT). Animals were then incubated with an appropriate antibody (1:3000 Roche anti-DIG-AP 11093274910, 1:1000 Roche anti-DIG-POD 11207733910 or 1:1000 Roche anti-FITC-POD 11426346910 in blocking solution) at 4°C overnight. Subsequent washes and tyramide development were performed as previously described. After development, animals were mounted in ScaleA2 [48] (4M urea, 20% glycerol, 0.1% Triton X-100, 2.5% DABCO). Whole-mount *in situ* hybridizations were either imaged on a Leica M165F with a DFC7000T camera or on a Zeiss 710 confocal microscope, and images were processed in Fiji [49]. *smedwi-1* stem cells were quantified manually and statistical analysis was carried out using JMP Pro statistical analysis software.

### Phosphohistone H3 labeling

Animals were stained with phosphohistone H3 following *in situ* hybridizations. After inactivation of peroxidase with 200mM sodium azide in PBSTx for 1 hour at RT, animals were rinsed in >6 PBSTx washes. Then, animals were incubated in anti-phosphohistone H3 (Ser10) antibody (Abcam, Cambridge, MA Ab32107) at a concentration of 1:1000 in blocking solution for 48 hours at 4°C. Primary was washed off with PBSTx followed by incubation with a goat anti-rabbit-HRP secondary antibody (Thermo Fisher) at a concentration of 1:2000 in PBSTx overnight at 4°C. Antibody was washed off with PBSTx and samples were pre-incubated in fluorescein tyramide (1:5000 in PBSTx) for 10 minutes and then developed with 0.005% H_2_O_2_ in PBSTx for 10 minutes at room temperature. After development, samples were rinsed in PBSTx and counterstained with DAPI (Thermo Scientific) diluted 1:5000 in PBSTx before mounting in ScaleA2. Animals were imaged on a Zeiss 710 confocal microscope and images were processed in Fiji [49].

### Flow cytometry and Annexin V staining

Flow cytometry of Hoechst-stained cells was conducted as previously described [11,26] with minor modifications. Sixty animals per group were placed in 0.065% NAC for 1 minute, followed by rinsing with Montjuïc salt solution. Pre-pharyngeal regions from all animals were dissected and rinsed in CMFB (CMF+0.5% BSA). These fragments were pooled and dissociated using 1:100 Liberase™ (2.5mg/ml, Roche 5401135001) in CMFB at 30°C with gentle agitation at 300 rpm for 30 minutes on an Eppendorf ThermoMixer™. Samples were gently triturated every 5 minutes to aid dissociation. Dissociated cells were then diluted with equal volume of CMFB and pelleted by centrifugation (500g, 5 minutes, RT). Pelleted cells were diluted in 1ml of CMFB and strained using a 30µm cell strainer (BD 340627). Strained cells were counted using an automated cell counter (Beckmann Coulter particle counter) and 2.8×10^6^ cells/group were stained with 5µg/ml Hoechst 33342 (ThermoFisher H3570) in CFMB for 70 mins in the dark with gentle agitation. Cells were subsequently pelleted and Hoechst solution was replaced with 0.5µl of Annexin V-APC (Thermo Fisher A35110) in 100µl freshly made 1X Annexin V buffer from a 10X stock solution (0.1M HEPES pH 7.4, 1.4M NaCl, and 25 mM CaCl_2_). After staining for 15 minutes at RT, 400µl of 1X Annexin V buffer with 1µg/ml Propidium Iodide (Sigma P4170) was added to each tube. Cells were strained again immediately before analysis on a BD FACS AriaII cell sorter. Flow cytometry data was analyzed in FlowJo (TreeStar, Ashland, OR).

### TUNEL staining

TUNEL staining protocol was adapted from [2]. Animals fixed as above were rehydrated and bleached overnight with 6% hydrogen peroxide in PBSTx. Bleached animals were rinsed in PBSTx, permeabilized with Proteinase K (2µg/ml in PBSTx) and post-fixed in 4% formaldehyde for 10 minutes in PBSTx. Animals were rinsed with PBS and transferred to 1.5mL microcentrifuge tubes with a maximum of 5 animals/tube. PBS was replaced with 25µl reaction mix: 3 parts ApopTag TdT enzyme mix, 7 parts ApopTag reaction buffer (Millipore S7107) and incubated overnight at 37°C. Animals were washed with in PBSTx, and blocked for 2 hours (0.5% Roche Western Blocking Reagent and 5% inactivated horse serum in PBSTx) at room temperature. Animals were incubated in antibody (1:1000 anti-DIG-POD in blocking solution) at 4°C for 24 hours. Antibody was washed off with PBSTx and developed using Cy3-tyramide (1:1000 in PBSTx, with 0.005% H_2_O_2_) for 10 minutes. After development, samples were counterstained with DAPI (Thermo Scientific) 1:5000 in PBSTx and mounted in ScaleA2, and imaged on a Zeiss 710 confocal microscope. Images were processed in Fiji.

### Cycloheximide treatment

Cycloheximide (VWR 97064-722) was administered at a concentration of 4µM in planaria water containing a final concentration of 1.5% DMSO. All animals were soaked in cycloheximide for a total of 72 hours. Unirradiated animals were transferred to planaria water 24 hours prior to fixation. Animals subjected to radiation were soaked 48 hours prior to exposure and maintained in cycloheximide for 24 hours after radiation, then washed and kept in planaria water for another 24 hours before fixing.

### TUNEL-FISH

Animals were transferred to 1.5ml eppendorf tubes and incubated in 10% NAC in PBS for 5 minutes, then fixed in 4% paraformaldehyde in PBSTx for 20 minutes at room temperature. After fixation, animals were rinsed twice in PBSTx and permeabilized with proteinase K (20µg/ml) at 37°C for 10 minutes. Proteinase K was replaced with pre-warmed reduction solution as above and then fixed in 4% paraformaldehyde for 10 minutes. After 2 washes, animals were bleached in 6% H_2_O_2_ in 100% methanol overnight. Animals were then rinsed in 1X PBS, and 6 animals/tube were incubated in Tdt reaction mix (as above), for 4 hours. Tdt reaction mix was then washed 4x times with PBSTx over 1 hour followed by hybridization with riboprobe as above. The riboprobe was developed using Fluorescein-tyramide as previously described. This was followed by inactivation of peroxidase as above. After washing, animals were incubated overnight at 4°C with anti-DIG-POD (1:1000) in blocking solution. After washing, TUNEL was developed using Cy3 tyramide as above. Animals were then stained with DAPI, mounted and imaged as above.

### RNAi

RNAi was carried out as previously described [50]. Briefly, double stranded RNA (dsRNA) was synthesized in vitro using PCR products of genes mentioned below as a template and mixed with a 4:1 liver:water paste. Animals were fed a total of 4µg of dsRNA per 10µl of food every 3 days for a total period of 12 days. Radiation and amputation was carried out 7 days after the last feed and animals were fixed 48 hours later.

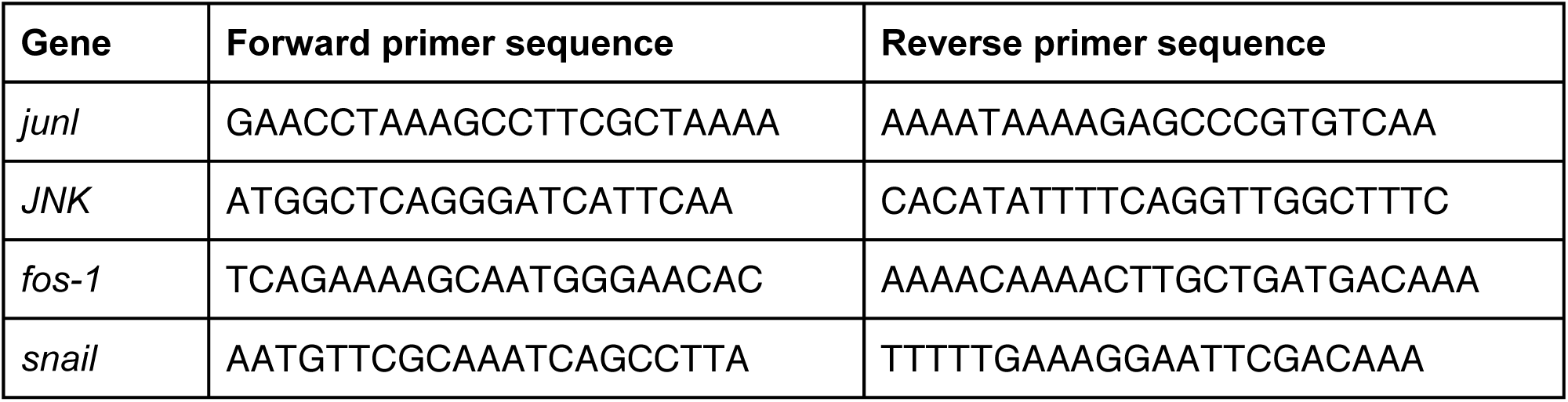

## REFERENCES

1. Wagner, D.E., Wang, I.E., and Reddien, P.W. (2011). Clonogenic neoblasts are pluripotent adult stem cells that underlie planarian regeneration. Science 332, 811–816.

2. Pellettieri, J., Fitzgerald, P., Watanabe, S., Mancuso, J., Green, D.R., and Sánchez Alvarado, A. (2010). Cell death and tissue remodeling in planarian regeneration. Dev. Biol. 338, 76–85.

3. Newmark, P.A., and Sánchez Alvarado, A. (2002). Not your father’s planarian: a classic model enters the era of functional genomics. Nat. Rev. Genet. 3, 210–219.

4. Rink, J.C. (2013). Stem cell systems and regeneration in planaria. Dev. Genes Evol. 223, 67–84.

5. Adler, C.E., and Sánchez Alvarado, A. (2015). Types or states? Cellular dynamics and regenerative potential. Trends Cell Biol. 25, 687–696.

6. Wenemoser, D., and Reddien, P.W. (2010). Planarian regeneration involves distinct stem cell responses to wounds and tissue absence. Dev. Biol. 344, 979–991.

7. Baguñà, J. (1976). Mitosis in the intact and regenerating planarian *Dugesia mediterranea* n.sp. II. Mitotic studies during regeneration, and a possible mechanism of blastema formation. J. Exp. Zool. 195, 65–79.

8. Wurtzel, O., Cote, L.E., Poirier, A., Satija, R., Regev, A., and Reddien, P.W. (2015). A generic and cell-type-specific wound response precedes regeneration in planarians. Dev. Cell 35, 632–645.

9. Sandmann, T., Vogg, M.C., Owlarn, S., Boutros, M., and Bartscherer, K. (2011). The head-regeneration transcriptome of the planarian *Schmidtea mediterranea*. Genome Biol. 12, R76.

10. Bardeen, C.R., and Baetjer, F.H. (1904). The inhibitive action of the Roentgen rays on regeneration in planarians. J. Exp. Zool.

11. Reddien, P.W., Oviedo, N.J., Jennings, J.R., Jenkin, J.C., and Sánchez Alvarado, A. (2005). SMEDWI-2 is a PIWI-like protein that regulates planarian stem cells. Science 310, 1327–1330.

12. Lei, K., Thi-Kim Vu, H., Mohan, R.D., McKinney, S.A., Seidel, C.W., Alexander, R., Gotting, K., Workman, J.L., and Sánchez Alvarado, A. (2016). Egf signaling directs neoblast repopulation by regulating asymmetric cell division in planarians. Dev. Cell 38, 413–429.

13. Scimone, M.L., Meisel, J., and Reddien, P.W. (2010). The Mi-2-like Smed-CHD4 gene is required for stem cell differentiation in the planarian *Schmidtea mediterranea*. Development 137, 1231–1241.

14. Eisenhoffer, G.T., Kang, H., and Sánchez Alvarado, A. (2008). Molecular analysis of stem cells and their descendants during cell turnover and regeneration in the planarian Schmidtea mediterranea. Cell Stem Cell 3, 327–339.

15. Labbé, R.M., Irimia, M., Currie, K.W., Lin, A., Zhu, S.J., Brown, D.D.R., Ross, E.J., Voisin, V., Bader, G.D., Blencowe, B.J., et al. (2012). A Comparative Transcriptomic Analysis Reveals Conserved Features of Stem Cell Pluripotency in Planarians and Mammals. Stem Cells 30, 1734–1745.

16. Solana, J., Kao, D., Mihaylova, Y., Jaber-Hijazi, F., Malla, S., Wilson, R., and Aboobaker, A. (2012). Defining the molecular profile of planarian pluripotent stem cells using a combinatorial RNAseq, RNA interference and irradiation approach. Genome Biol. 13, R19.

17. van Wolfswinkel, J.C., Wagner, D.E., and Reddien, P.W. (2014). Single-cell analysis reveals functionally distinct classes within the planarian stem cell compartment. Cell Stem Cell 15, 326–339.

18. Guo, T., Peters, A.H.F.M., and Newmark, P.A. (2006). A Bruno-like gene is required for stem cell maintenance in planarians. Dev. Cell 11, 159–169.

19. Wagner, D.E., Ho, J.J., and Reddien, P.W. (2012). Genetic regulators of a pluripotent adult stem cell system in planarians identified by RNAi and clonal analysis. Cell Stem Cell 10, 299–311.

20. Onal, P., Grün, D., Adamidi, C., Rybak, A., Solana, J., Mastrobuoni, G., Wang, Y., Rahn, H.-P., Chen, W., Kempa, S., et al. (2012). Gene expression of pluripotency determinants is conserved between mammalian and planarian stem cells. EMBO J. 31, 2755–2769.

21. Zeng, A., Li, H., Guo, L., Gao, X., McKinney, S., Wang, Y., Yu, Z., Park, J., Semerad, C., Ross, E., et al. (2018). Prospectively isolated tetraspanin+ neoblasts are adult pluripotent stem cells underlying planaria regeneration. Cell 173, 1593–1608.e20.

22. Tu, K.C., Cheng, L.-C., T K Vu, H., Lange, J.J., McKinney, S.A., Seidel, C.W., and Sánchez Alvarado, A. (2015). Egr-5 is a post-mitotic regulator of planarian epidermal differentiation. eLife 4, e10501.

23. Saló, E., and Baguñà, J. (1984). Regeneration and pattern formation in planarians. I. The pattern of mitosis in anterior and posterior regeneration in *Dugesia (G) tigrina*, and a new proposal for blastema formation. J. Embryol. Exp. Morphol. 83, 63–80.

24. Oviedo, N.J., Newmark, P.A., and Sánchez Alvarado, A. (2003). Allometric scaling and proportion regulation in the freshwater planarian *Schmidtea mediterranea*. Dev. Dyn. 226, 326–333.

25. Lapan, S.W., and Reddien, P.W. (2012). Transcriptome analysis of the planarian eye identifies *ovo* as a specific regulator of eye regeneration. Cell Rep. 2, 294–307.

26. Hayashi, T., Asami, M., Higuchi, S., Shibata, N., and Agata, K. (2006). Isolation of planarian X-ray-sensitive stem cells by fluorescence-activated cell sorting. Dev. Growth Differ. 48, 371–380.

27. Bender, C.E., Fitzgerald, P., Tait, S.W.G., Llambi, F., McStay, G.P., Tupper, D.O., Pellettieri, J., Sánchez Alvarado, A., Salvesen, G.S., and Green, D.R. (2012). Mitochondrial pathway of apoptosis is ancestral in metazoans. Proc. Natl. Acad. Sci. U. S. A. 109, 4904–4909.

28. Crowley, L.C., Marfell, B.J., Scott, A.P., and Waterhouse, N.J. (2016). Quantitation of Apoptosis and Necrosis by Annexin V Binding, Propidium Iodide Uptake, and Flow Cytometry. Cold Spring Harb. Protoc. *2016*. Available at: http://dx.doi.org/10.1101/pdb.prot087288.

29. Wenemoser, D., Lapan, S.W., Wilkinson, A.W., Bell, G.W., and Reddien, P.W. (2012). A molecular wound response program associated with regeneration initiation in planarians. Genes Dev. 26, 988–1002.

30. Guedelhoefer, O.C., 4th, and Sánchez Alvarado, A. (2012). Amputation induces stem cell mobilization to sites of injury during planarian regeneration. Development 139, 3510–3520.

31. Almuedo-Castillo, M., Crespo-Yanez, X., Crespo, X., Seebeck, F., Bartscherer, K., Salò, E., and Adell, T. (2014). JNK controls the onset of mitosis in planarian stem cells and triggers apoptotic cell death required for regeneration and remodeling. PLoS Genet. 10, e1004400.

32. Tejada-Romero, B., Carter, J.-M., Mihaylova, Y., Neumann, B., and Aboobaker, A.A. (2015). JNK signalling is necessary for a Wnt- and stem cell-dependent regeneration programme. Development 142, 2413–2424.

33. Abnave, P., Aboukhatwa, E., Kosaka, N., Thompson, J., Hill, M.A., and Aboobaker, A.A. (2017). Epithelial-mesenchymal transition transcription factors control pluripotent adult stem cell migration in vivo in planarians. Development 144, 3440–3453.

34. Brock, C.K., Wallin, S.T., Ruiz, O.E., Samms, K.M., Mandal, A., Sumner, E.A., and Eisenhoffer, G.T. (2019). Stem cell proliferation is induced by apoptotic bodies from dying cells during epithelial tissue maintenance. Nat. Commun. 10, 1044.

35. Ryoo, H.D., Gorenc, T., and Steller, H. (2004). Apoptotic cells can induce compensatory cell proliferation through the JNK and the Wingless signaling pathways. Dev. Cell 7, 491–501.

36. Huh, J.R., Guo, M., and Hay, B.A. (2004). Compensatory proliferation induced by cell death in the *Drosophila* wing disc requires activity of the apical cell death caspase Dronc in a nonapoptotic role. Curr. Biol. 14, 1262–1266.

37. Fan, Y., and Bergmann, A. (2008). Apoptosis-induced compensatory proliferation. The Cell is dead. Long live the Cell! Trends Cell Biol. 18, 467–473.

38. Xing, Y., Su, T.T., and Ruohola-Baker, H. (2015). Tie-mediated signal from apoptotic cells protects stem cells in *Drosophila melanogaster*. Nat. Commun. 6, 7058.

39. Bilak, A., Uyetake, L., and Su, T.T. (2014). Dying cells protect survivors from radiation-induced cell death in *Drosophila*. PLoS Genet. 10, e1004220.

40. Verghese, S., and Su, T.T. (2018). Ionizing radiation induces stem cell-like properties in a caspase-dependent manner in *Drosophila*. PLoS Genet. 14, e1007659.

41. Adams, K.W., and Cooper, G.M. (2007). Rapid turnover of mcl-1 couples translation to cell survival and apoptosis. J. Biol. Chem. 282, 6192–6200.

42. Ma, M., Zhao, H., Zhao, H., Binari, R., Perrimon, N., and Li, Z. (2016). Wildtype adult stem cells, unlike tumor cells, are resistant to cellular damages in *Drosophila*. Dev. Biol. 411, 207–216.

43. Li, F., Huang, Q., Chen, J., Peng, Y., Roop, D.R., Bedford, J.S., and Li, C.-Y. (2010). Apoptotic cells activate the “phoenix rising” pathway to promote wound healing and tissue regeneration. Sci. Signal. 3, ra13.

44. Chera, S., Ghila, L., Dobretz, K., Wenger, Y., Bauer, C., Buzgariu, W., Martinou, J.-C., and Galliot, B. (2009). Apoptotic cells provide an unexpected source of Wnt3 signaling to drive hydra head regeneration. Dev. Cell 17, 279–289.

45. Newmark, P.A., and Sánchez Alvarado, A. (2000). Bromodeoxyuridine specifically labels the regenerative stem cells of planarians. Dev. Biol. 220, 142–153.

46. Pearson, B.J., Eisenhoffer, G.T., Gurley, K.A., Rink, J.C., Miller, D.E., and Sánchez Alvarado, A. (2009). Formaldehyde-based whole-mount *in situ* hybridization method for planarians. Dev. Dyn. 238, 443–450.

47. King, R.S., and Newmark, P.A. (2013). In situ hybridization protocol for enhanced detection of gene expression in the planarian *Schmidtea mediterranea*. BMC Dev. Biol. 13.

48. Hama, H., Kurokawa, H., Kawano, H., Ando, R., Shimogori, T., Noda, H., Fukami, K., Sakaue-Sawano, A., and Miyawaki, A. (2011). Scale: a chemical approach for fluorescence imaging and reconstruction of transparent mouse brain. Nat. Neurosci. 14, 1481–1488.

49. Schindelin, J., Arganda-Carreras, I., Frise, E., Kaynig, V., Longair, M., Pietzsch, T., Preibisch, S., Rueden, C., Saalfeld, S., Schmid, B., et al. (2012). Fiji: an open-source platform for biological-image analysis. Nat. Methods 9, 676–682.

50. Rouhana, L., Weiss, J.A., Forsthoefel, D.J., Lee, H., King, R.S., Inoue, T., Shibata, N., Agata, K., and Newmark, P.A. (2013). RNA interference by feeding in vitro-synthesized double-stranded RNA to planarians: methodology and dynamics. Dev. Dyn. 242, 718–730.

