## Supplementary figures and images for "Injury stimulates stem cells to resist radiation-induced apoptosis"

### Supplementary Figure 1

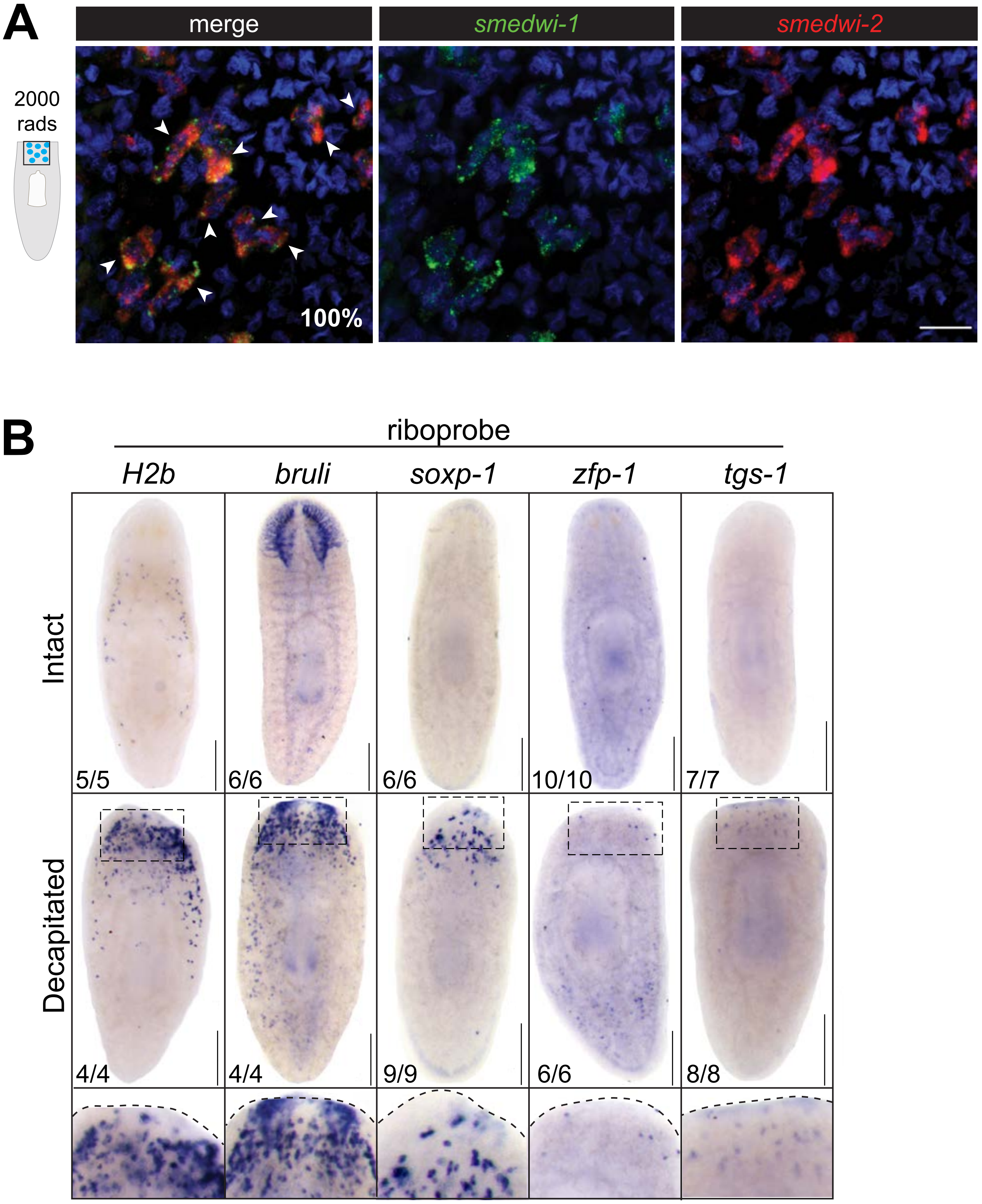

### Supplementary Figure 2

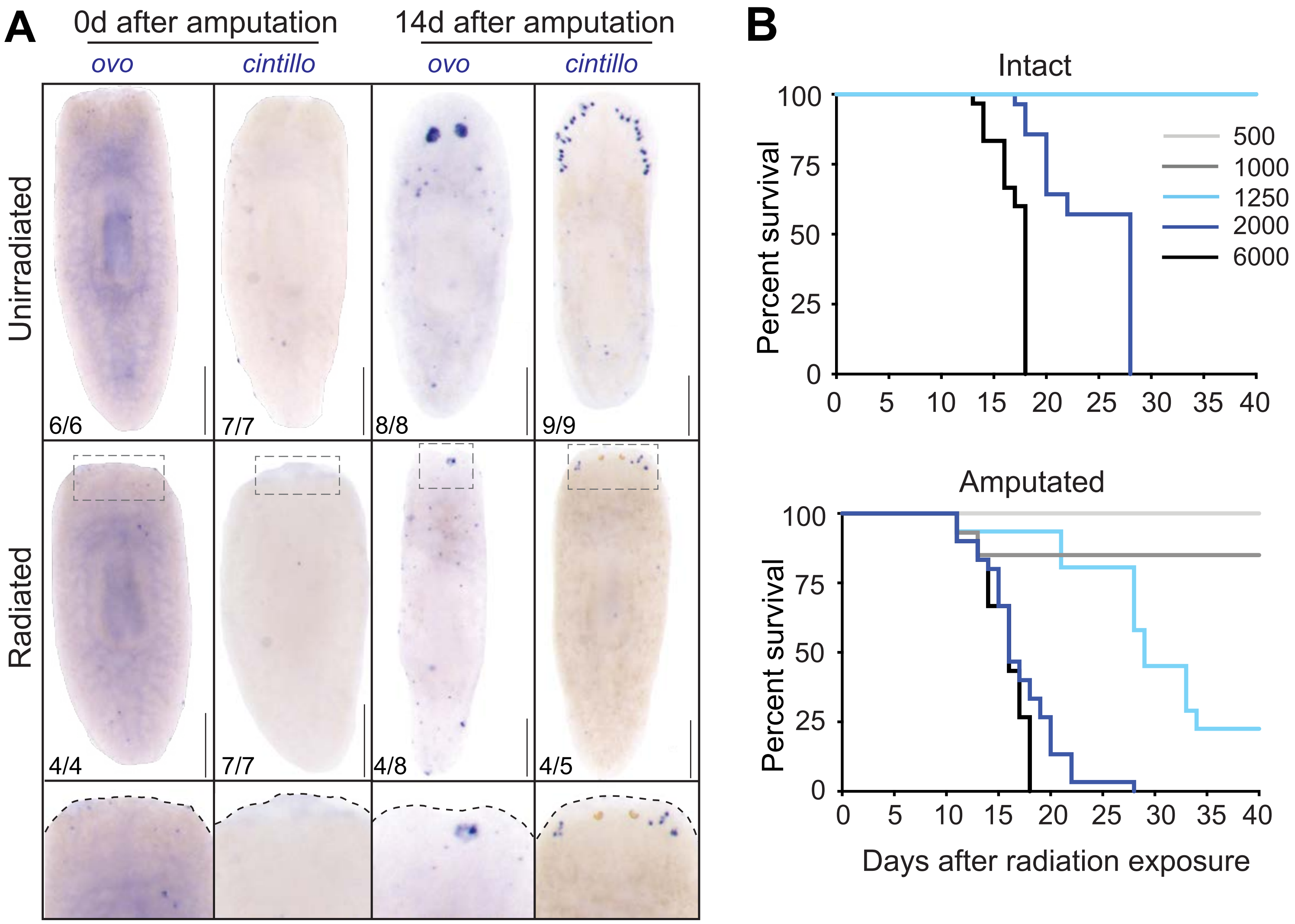

### Supplementary Figure 3

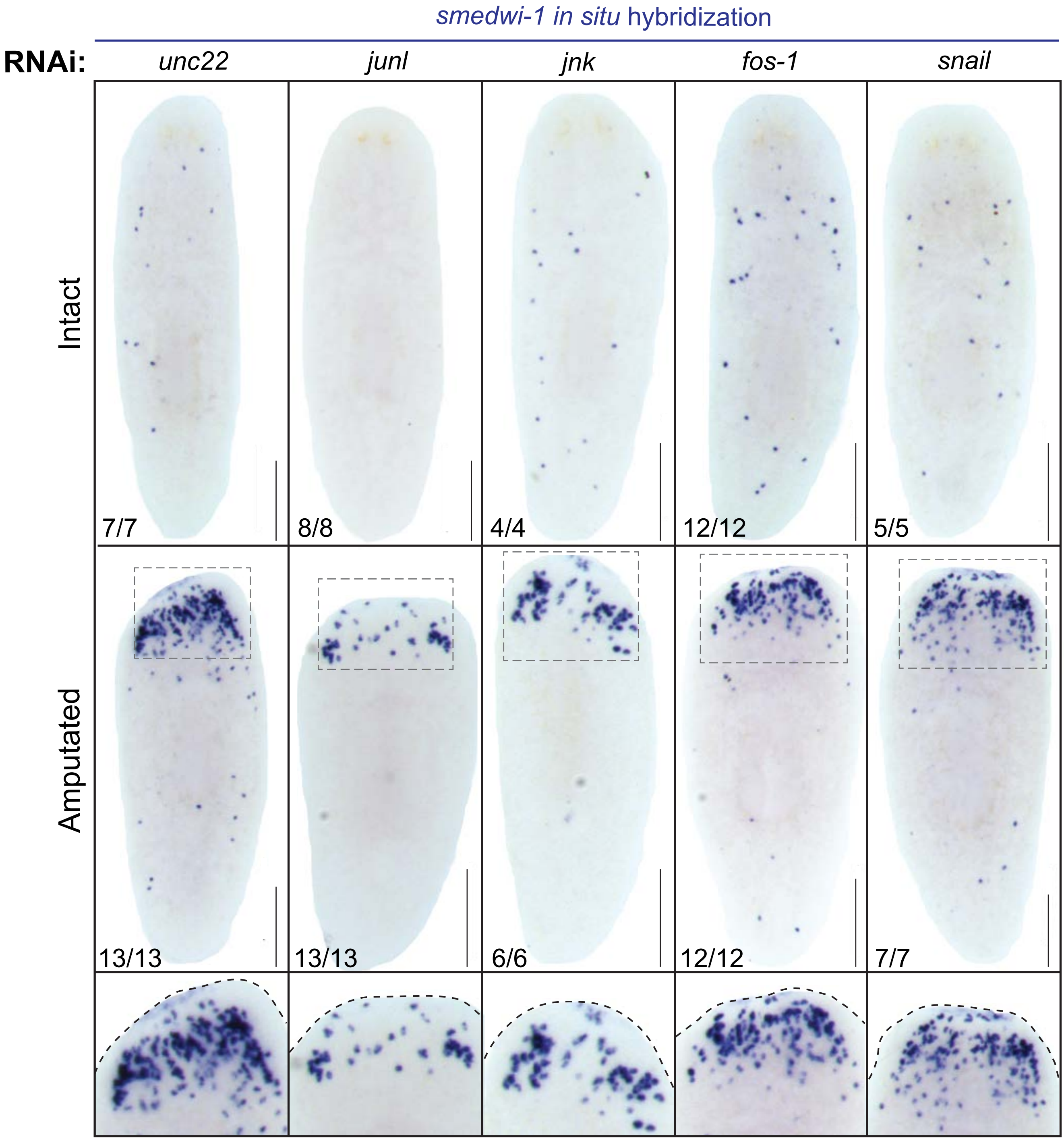
